# Metagenomic evidence for a polymicrobial signature of sepsis

**DOI:** 10.1101/2020.04.07.028837

**Authors:** Cedric Chih Shen Tan, Mislav Acman, Lucy van Dorp, Francois Balloux

**Author notes:** Co-last authors (Lucy van Dorp, Francois Balloux).

## Abstract

Our understanding of the host component of sepsis has made significant progress. However, detailed study of the microorganisms causing sepsis, either as single pathogens or microbial assemblages, has received far less attention. Metagenomic data offer opportunities to characterise the microbial communities found in septic and healthy individuals. In this study we apply gradient-boosted tree classifiers and a novel computational decontamination technique built upon SHapley Additive exPlanations (SHAP) to identify microbial hallmarks which discriminate blood metagenomic samples of septic patients from that of healthy individuals. Classifiers had high performance when using the read assignments to microbial genera (AUROC = 0.995), including after removal of species ‘confirmed’ as the cause of sepsis through clinical testing (AUROC = 0.915). Models trained on single genera were inferior to those employing a polymicrobial model and we identified multiple co-occurring bacterial genera absent from healthy controls.

**Importance:** While prevailing diagnostic paradigms seek to identify single pathogens, our results point to the involvement of a polymicrobial community in sepsis. We demonstrate the importance of the microbial component in characterising sepsis, which may offer new biological insights into the aetiology of sepsis and allow the development of clinical diagnostic or even prognostic tools.

## Introduction

Sepsis poses a significant challenge to public health and was listed as a global health priority by the World Health Organisation (WHO) in 2017. In the same year, 48.9 million cases of sepsis and 11 million deaths were recorded worldwide [1] having a particular impact in low and lower-middle income countries [2].

Current research efforts have predominately focused on understanding the host’s response to sepsis. Indeed, all contemporary definitions of sepsis focus on the host’s response and resulting systemic complications. The 1991 Sepsis-1 definition described sepsis as a systemic inflammatory response syndrome (SIRS) caused by infection, with patients being diagnosed with sepsis if they fulfil at least two SIRS criteria and have a clinically confirmed infection [3]. The 2001 Sepsis-2 definition then expanded the scope of SIRS to include more symptoms [4]. More recently, the 2016 Sepsis-3 definition sought to differentiate between mild and severe cases of dysregulated host responses, describing sepsis as a life-threatening organ dysfunction as a result of infection [5]. Significant progress has been made in understanding how dysregulation occurs [6] and the long-term impacts of sepsis [7,8]. Additionally, early-warning tools have been developed based on patient health-care records [9–11] and clinical checklists [12,13]. However, the focus on the host component of sepsis may overlook the important role of microbial composition in the pathogenesis of the disease.

Due to the severity of sepsis, current practice considers identification of a single pathogen sufficient to warrant a diagnosis, without consideration of other, potentially relevant, species in the bloodstream. Upon diagnosis, infections are rapidly treated with broad spectrum antibiotics. However, blood cultures, the current recommended method of diagnosis before antimicrobial treatment [14], are known to yield false negatives due to certain microorganisms failing to grow in culture [15], particularly in samples with low microbial loads [16]. Culture-based methods, while useful in a clinical context, may therefore under-estimate the true number of causative pathogens infecting septic patients.

Sepsis is a highly heterogeneous disease which consists of both a host component and a microbial component. While the former has been widely studied, the latter appears to represent a largely untapped source of information that could further advance our understanding of sepsis. Several diseases manifest as a result of interactions in a polymicrobial community. For example, microbial interactions in lung, urinary tract and wound infections are all known to contribute to differing disease outcomes (reviewed by Tay *et al.* [17]). These findings suggest that the microbial component of sepsis may also be crucial to understanding its pathogenesis.

Current technologies to investigate the presence of polymicrobial communities have some major limitations. As noted previously, culture-based methods have a high false negative rate. Further, without knowledge of the range of microorganisms that infect blood, co-culture experiments to study microbial interactions prove difficult. For polymerase chain reaction-based technologies, the use of species-specific primers (e.g. SeptiFast [18]) necessitates *a priori* knowledge of microbial sequences endogenous to septic blood. Lastly, metagenomic sequencing is ubiquitously prone to environmental contamination. This can include DNA from viable cells introduced during sample collection, sample processing, or DNA present in laboratory reagents [19–21]; the so called ‘kitome’. As such, it can be difficult to determine which microorganisms are truly endogenous to the sample, and at what abundance.

In this study, we sought to expand our understanding of the full microbial component of sepsis. Multiple statistical and state-of-the-art machine learning techniques were applied to metagenomic sequencing data published by Blauwkamp *et al*. [22] (henceforth Karius study) from 117 sepsis patients and 170 healthy individuals. To circumvent the problem of potential contamination in metagenomic data, we developed and applied a novel computational contamination reduction technique. We also externally validated our findings using external hold-out datasets comprising three other independent sepsis cohorts. Taken together, our results provide strong evidence for a polymicrobial signature of sepsis and the utility of metagenomic sequencing for the investigation of blood-borne infections.

## Results

### Metagenomic sequencing can be used to discriminate septic from healthy samples

We first assessed the suitability of taxonomic assignments for discriminating between septic and healthy blood metagenomic samples. Gradient-boosted tree classifiers were trained and evaluated using data matrices generated via *Kraken 2* taxonomic assignment, with samples represented in rows and taxa in columns *(i.e.* features). Each element in the matrices represented the total number of reads assigned to each taxon, which we loosely refer to as ‘abundance’. The set of taxa used in each analysis will henceforth be referred to as the ‘feature space’. Models were first trained and evaluated using 117 septic patients and 170 healthy individuals in the Karius study (Table 1). To determine if our findings were applicable beyond the Karius dataset, we pooled the Karius dataset with metagenomic information from three other independent sepsis cohorts [23–25]. The final pooled dataset contains sequence data from multiple sources, sepsis definitions and sequencing techniques (Table 1). We will henceforth refer to individual datasets by their dataset alias as shown in Table 1.

**Table 1.** Summary of metagenomic datasets. Sample sizes indicated here are those after all quality control steps have been applied.

| Study | Dataset alias | Accession | Sepsis definition | Sequencing technique | Sample size |  |
| --- | --- | --- | --- | --- | --- | --- |
|  |  |  |  |  | Septic | Healthy |
| Grumaz *et al.* (2019) | Grumaz-19 | PRJEB21872<br>PRJEB30958 | Sepsis-2 | Shotgun | 50 | - |
| Grumaz *et al.* (2016) | Grumaz-16 | PRJEB13247 | Sepsis-2 | Shotgun | 7 | 15 |
| Gosiewski *et al.* (2017) | Gosiewski-17 | Requested from authors | Sepsis-1 | 16S (paired-end) | 56 | 23 |
| Blauwkamp *et al.* (2019) | Karius | PRJNA507824 | Sepsis-1 | Shotgun | 117 | 170 |

The performance of all classifiers is summarised in Table 2. Using the raw feature space, parsed from the *Kraken 2* taxonomic assignments, classifiers had a very high classification performance *(Karius-Neat* model; AUROC = 0.995) in discriminating sepsis from healthy samples based on microbial content alone. This was similarly observed when using the pooled dataset *(Pooled-Neat* model; AUROC = 0.982).

**Table 2.** Summary of models trained. The prefix and suffix of each model name corresponds to the dataset and contamination reduction technique applied, respectively. *Neat, SD,* and *CR* refer to the feature spaces with no, Simple Decontamination, and SHAP Decontamination applied, respectively (see Methods). *Karius-Without* corresponds to the SHAP decontaminated feature space after claimed ‘confirmed’ pathogens are excluded. *Karius-Only* refers to the feature space containing only genera with ‘confirmed’ pathogens as features.

| No. of<br>Features | Feature Space | Model Performance |  |  |
| --- | --- | --- | --- | --- |
|  |  | Precision | Recall | AUROC |
| 1564 | *Karius-Neat* | 0.976 | 0.983 | 0.995 |
| 111 | *Karius-SD* | 0.896 | 0.787 | 0.942 |
| 25 | *Karius-CR* | 0.883 | 0.810 | 0.942 |
| 22 | *Karius-Without* | 0.803 | 0.727 | 0.915 |
| 22 | *Karius-Only* | 0.929 | 0.862 | 0.950 |
| 685 | *Pooled-Neat* | 0.950 | 0.939 | 0.982 |
| 21 | *Pooled-CR* | 0.870 | 0.796 | 0.904 |

### SHAP can be used to remove putative sequencing contaminants

Accurate characterisation of the microbial component of sepsis requires discrimination between a true biological signal and that arising from putative environmental contamination in metagenomes. We developed and applied a procedure to remove biologically irrelevant genera from the feature space, which we will refer to as SHAP Decontamination (CR; see Methods). Briefly, we leveraged SHapley Additive exPlanations (SHAP) – a state-of-the-art machine learning technique for interpreting ‘blackbox’ classifiers [26] – to determine how the read counts assigned to a genus *(i.e.* feature) influences model predictions. In doing so, we selectively removed putative contaminants from the feature spaces obtained from taxonomic classification.

To evaluate the effectiveness of this approach, we compared SHAP Decontamination to a simpler statistical method for the removal of putative pathogens, which we call Simple Decontamination (SD; see Methods). For the Karius dataset, application of SHAP Decontamination resulted in a pruned feature space of 25 genera while Simple Decontamination resulted in 111 genera. The resultant *Karius-CR* and *Karius-SD* feature spaces, respectively, shared 21 genera in common. Classifiers trained on either of the *Karius-CR* or *Karius-SD* feature space had similarly high performance (Table 2, *Karius-CR/SD;* AUROC = 0.942), despite the large reduction in the number of features. This suggests that computational decontamination efficiently removes redundancy in the metagenomic feature space. Furthermore, SHAP Decontamination appears to be more efficient as demonstrated by the equivalent classification performance, but higher number of removed putative contaminant genera than Simple Decontamination.

Separately, we observed that the *Karius-CR* model comprised almost all genera associated to sepsis at higher abundance. Additionally, genera such as *Sphingobium, Mesorhizobium* and *Ralstonia,* were highly important features in the *Karius-Neat* feature space (Fig. 1a), though not present in either the *Karius-SD* or *Karius-CR* feature space (Fig. 1b and c). These genera are likely to be contaminants since they contribute negatively to the predicted probability of sepsis at high abundance, and have been previously ascribed as common sequencing contaminants [19]. Of the 25 genera in the *Karius-CR* feature space, eight corresponded to genera containing clinically ‘confirmed’ pathogens (see Methods). Notably, *Escherichia* and *Enterobacter*, which are both ‘confirmed’ pathogens but also common contaminants [19], were retained in both decontaminated feature spaces. These findings collectively suggest that computational decontamination procedures were removing putative contaminants while selectively retaining biologically important genera.

**Figure 1.**
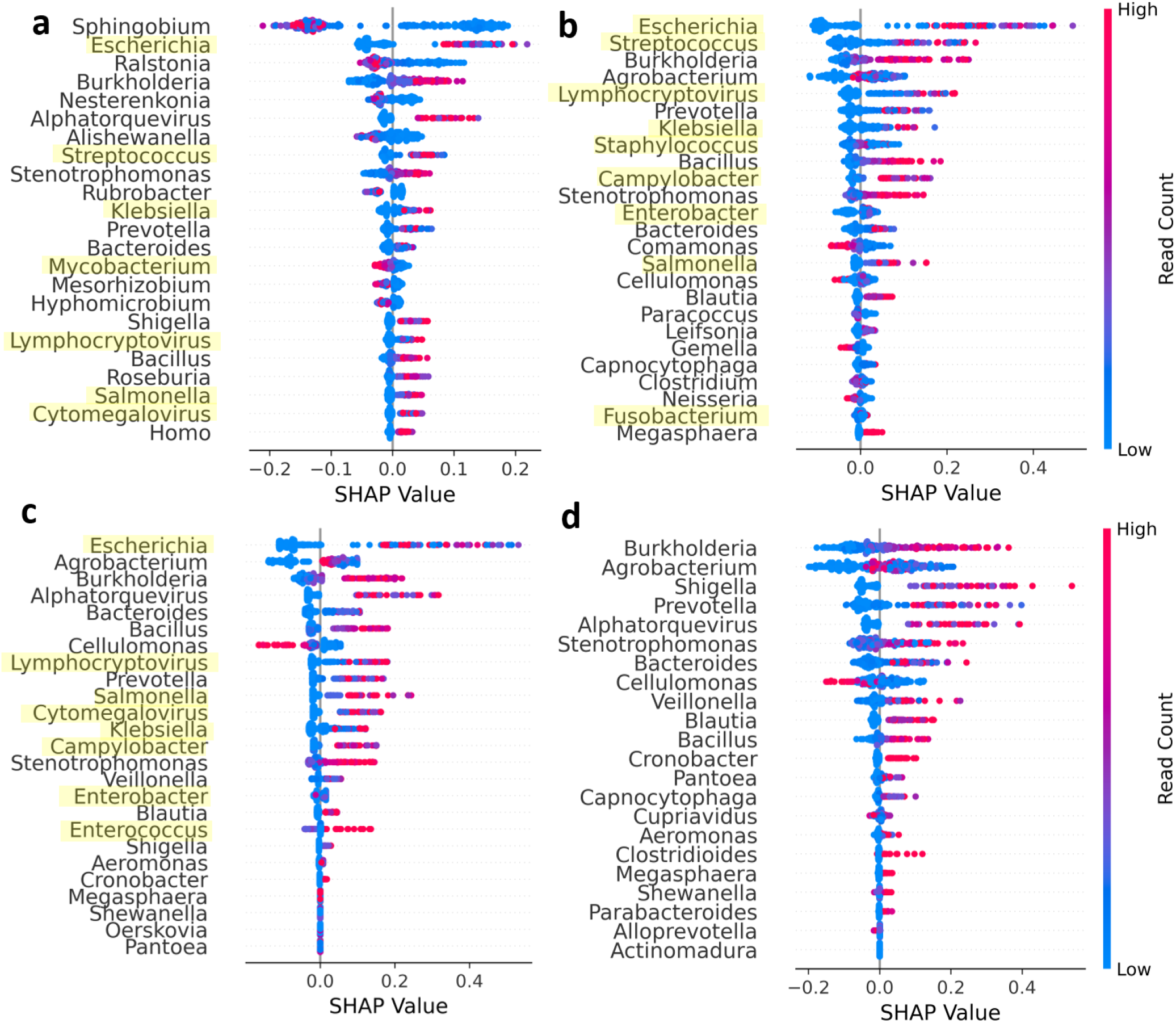
Model interpretation and performance. (a) Plot summarising the SHAP values across all samples for the most important features ranked by the mean absolute SHAP value (highest at the top) for *Karius-Neat*, (b) *Karius-*SD, (c) *Karius-CR* and (d) *Karius-Without* models. Each point represents a single sample. Points with similar SHAP values were stacked vertically for visualisation of point density and were coloured according to the magnitude of the feature values *(i.e.* read counts). Genera that contained ‘confirmed’ pathogens are highlighted in yellow.

### Evidence for a polymicrobial community

Having assessed the biological relevance of microbial predictors of sepsis, we provide several pieces of evidence supporting a polymicrobial model of sepsis; that is, that there are sets of microbial genera that delineate septic from healthy blood metagenomes, rather than just individual pathogens. Most notably, a classifier trained on the Karius dataset using the SHAP decontaminated feature space but with all genera containing clinically identified pathogens (henceforth ‘confirmed’ pathogens; see Methods) removed performed well *(Karius-Without* model; AUROC = 0.915) suggesting the presence of these species alone does not capture the full microbial signal of sepsis. Visualisation of the SHAP values for this model (Fig. 1d) confirmed that most genera had positive associations with sepsis at higher abundances. To test if any single features in the *Karius-Without* model were driving the high classification performance, we trained and evaluated multiple single-feature classifiers with each genus in the *Karius-Without* feature space. Additionally, we trained a classifier on genera containing ‘confirmed’ pathogens as features only (*Karius-Only*). Fig. 2 shows the performance of the multi-feature *Karius-Neat*, *Karius-Without* and *Karius-Only* models compared to single-feature models. All multi-feature models performed superior to those relying on single-feature models.

**Figure 2.**
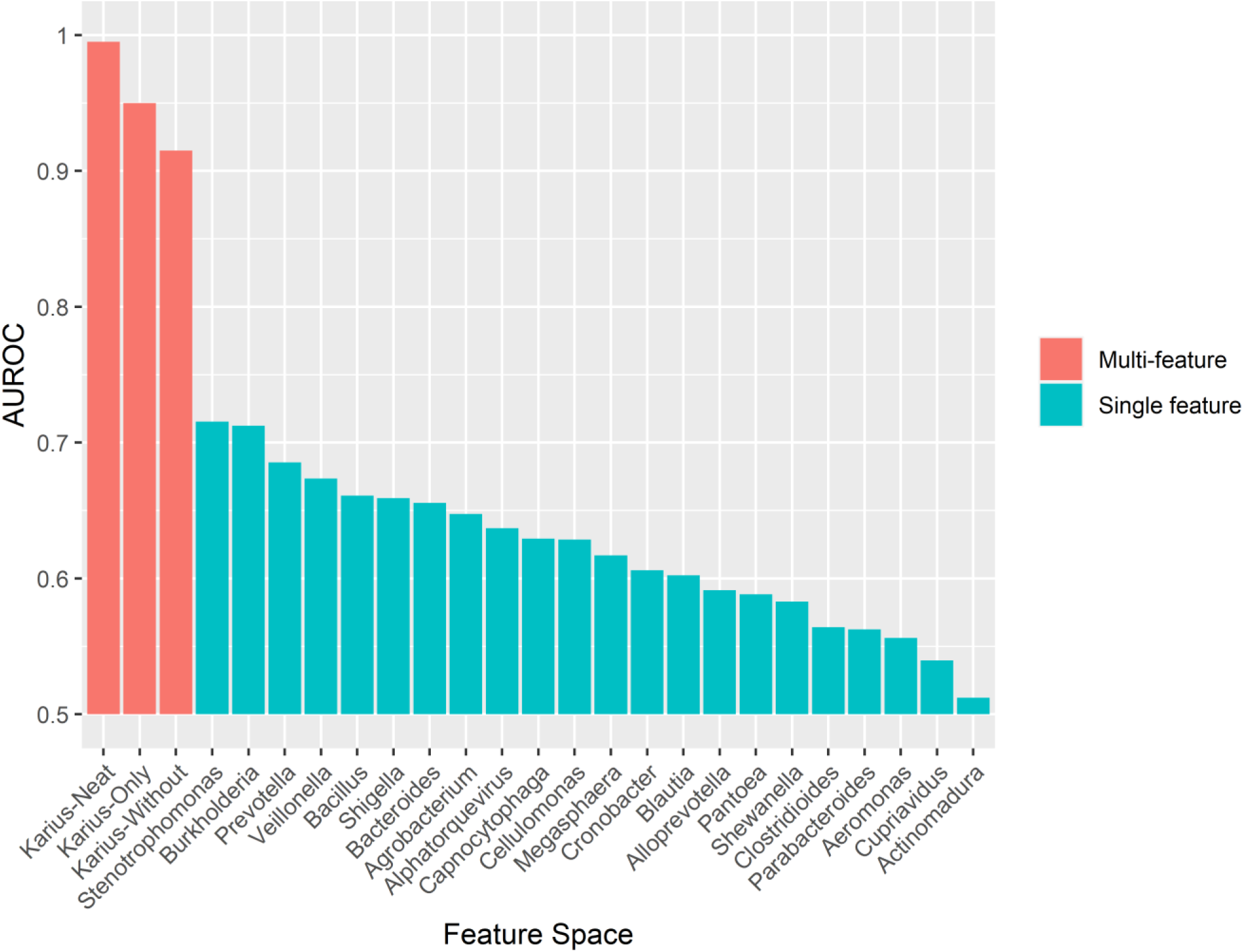
Comparison of performance (AUROC) for the multi-feature models *(Karius-Neat, Karius-Only, Karius-Without* feature space) and single-feature models (x-axis).

We then trained classifiers on the pooled dataset to determine if our results were unique to the Karius dataset or whether they were portable to other sepsis cohorts. Current metagenomics datasets are limited in their suitability for external validation due to the use of different sequencing technologies, differing sepsis definitions and small sample sizes. However, despite the pooled dataset comprising multiple data sources from different studies, the classifier still performed well (*Pooled-Neat* model, AUROC = 0.982; *Pooled-CR* model, AUROC = 0.904). This strongly suggests that there is a generalisable microbial signature which can be leveraged across metagenomic datasets.

To more formally test the generalisability of the observed polymicrobial signature, we trained classifiers on pooled data from two data sources while holding out data from the last source for testing (Fig. 3). Most notably, the classifier trained on shotgun metagenomic data and tested on 16S data as the holdout set (Gosiewski-17) did not perform well. However, after SHAP Decontamination, classification performance improved markedly. Interestingly, this performance increase was not observed when using the other datasets as holdout sets (Fig. 3). Indeed, the classifier trained on the feature space before SHAP Decontamination with the Sepsis-2, Grumaz-16 and Grumaz-19 datasets as holdout performed well, whereas that trained with the feature space after decontamination performed relatively worse. Additionally, holding out the Karius dataset resulted in poor classification performance both before and after SHAP Decontamination. A possible explanation for SHAP Decontamination lowering classification performance when Grumaz-16/19 is used as the test set is that septic cases recruited in these studies were based on different sepsis definitions which may involve a different set of pathogens and reflect different aetiologies. Separately, the poor performance observed when the Karius dataset is used as the test set can be attributed to the highly imbalanced training dataset (Fig. 3).

**Figure 3.**
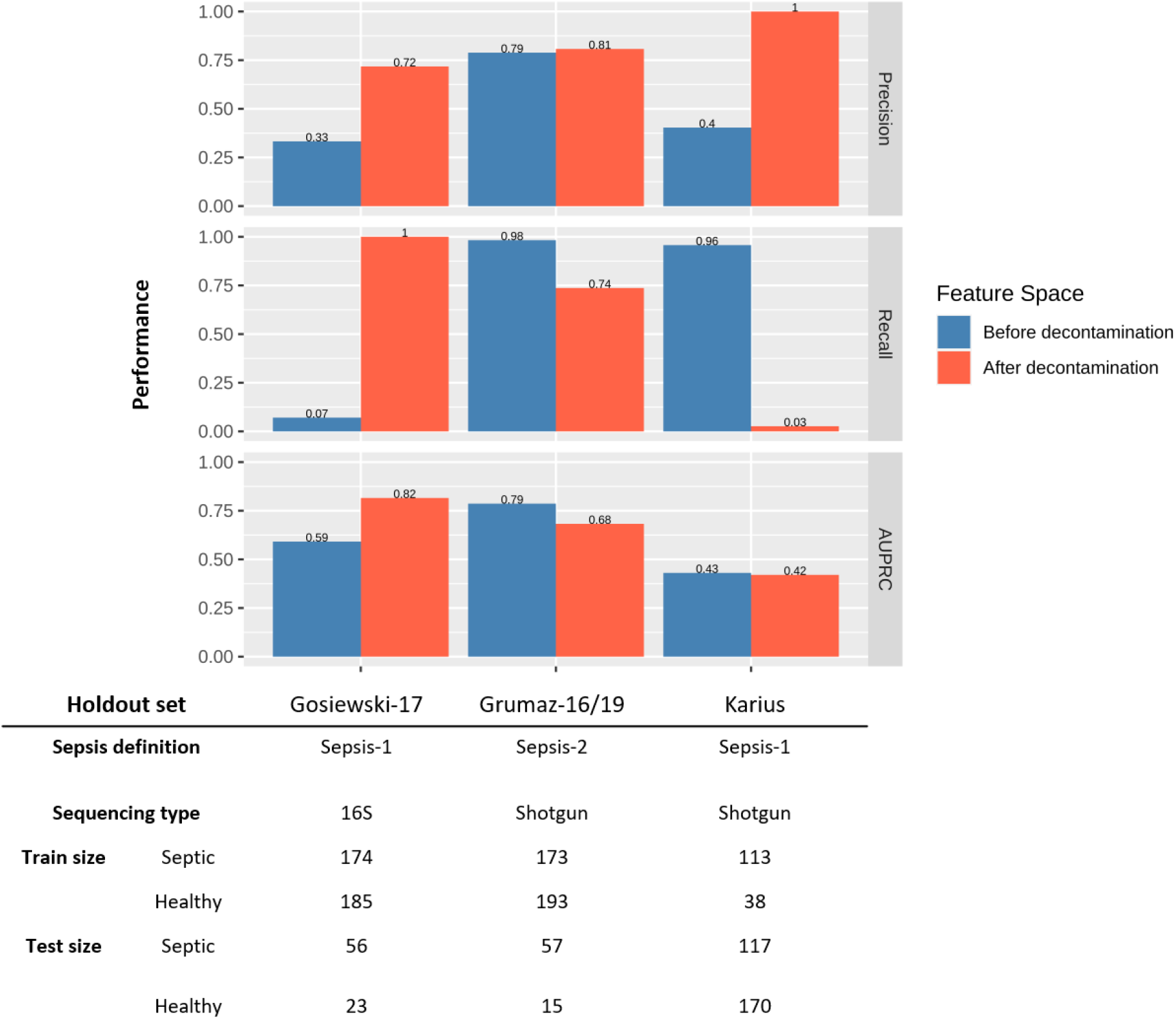
Performance of optimised classifiers tested on different holdout datasets before and after SHAP Decontamination. Grumaz-16 and Grumaz-19 were pooled to form a single test set.

Lastly, microbial co-occurrence networks were used to identify relationships between genera that were exclusive to samples from septic patients. Two genera are said to co-occur if an increase in the abundance of one is associated with an increase in the abundance of the other. The presence of such relationships would lend weight to the polymicrobial nature of sepsis infections. The *Karius-SD* feature space was used in this analysis to corroborate previous analyses using the *Karius-CR* feature space. Multiple co-occurrence relationships between genera were present in the corrected network including those containing 10 of the 22 ‘confirmed’ pathogens and 14 of the 25 genera in the *Karius-CR* feature space (Fig. 4). Interestingly, we detected a group of co-occurring genera associated to the oral cavity (Fig. 4), as suggested by the Human Oral Microbiome Database [27] (accessed 15^th^ July 2020) and the current literature [28–31]. This was also present in the corrected network when the *Pooled-SD* feature space was used as input (Fig. S1).

**Figure 4.**
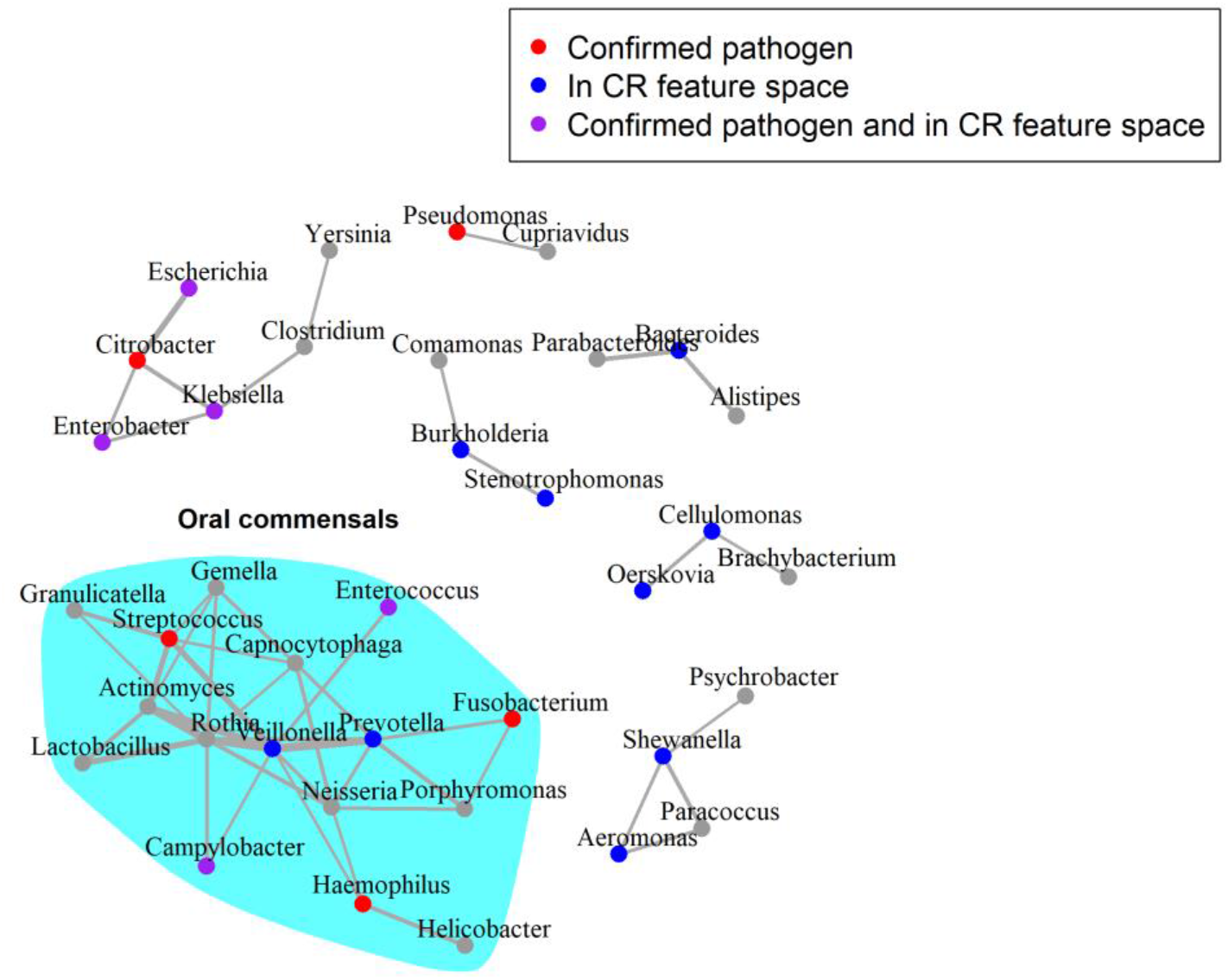
Corrected microbial co-occurrence network for genera assigned in sepsis metagenomes. Input data corresponds to the *Karius-SD* feature space. The edges in this network represent those in the septic network that were not present in the healthy network. The widths of edges are weighted by the strength of the *SparCC* correlations. Nodes are coloured as per the legend at top, with ‘confirmed’ pathogens those experimentally shown to be implicated in sepsis. The layout of the graph was generated using the Fruchterman-Reingold algorithm.

## Discussion

### The polymicrobial signature of sepsis

Our work demonstrates a clear polymicrobial signal in sepsis, where multiple, co-occuring, genera can be used to discriminate blood metagenomes of septic patients from that of healthy controls. The high performance of the *Karius-Without* model primarily highlights that genera containing ‘confirmed’ pathogens were very useful in delineating septic from healthy samples. More importantly, the *Karius-Without* model, which had these genera removed (*Karius-Without*) also performed well, suggesting that the abundance of microbial genera that were not amongst the ‘confirmed’ pathogens are also highly relevant to delineating septic from healthy samples. Furthermore, the single-feature models performed poorly, highlighting that no genus is solely responsible for the high classification performance of the *Karius-Without* model, further supporting the polymicrobial nature of sepsis infections.

We also show that the polymicrobial signal we detected is generalisable across datasets, first by nested cross-validation with all datasets pooled (*Pooled-CR* model) and then with holdout cross-validation using the Gosiewski-17 or Grumaz-16/19 datasets as test sets. The increased performance after SHAP Decontamination when holding out 16S data (Gosiewski-17) suggests that the retained set of genera allow a markedly more generalisable decision boundary to be learnt, even across sequencing techniques.

Additionally, the multiple co-occurrence relationships between genera detected suggest that there may be a distinct microbial community that tends to be present during sepsis infection. Although our networks were inferred computationally, published evidence supports possible synergies between some of the co-occurring genera we detected. For example, using fluorescence in-situ hybridisation, interspecies spatial associations were found between *Prevotella*, *Veillonella*, *Streptococcus*, *Gemella*, *Rothia* and *Actinomyces* [32], which were also the genera with the strongest correlations in the corrected sepsis network (Fig. 3). Separately, *Stenotrophomonas* and *Burkholderia* are known to play a collective role in the pathogenesis of cystic fibrosis [33]. Lastly, *Klebsiella pneumoniae* was found to be able to transmit extended spectrum beta-lactamase genes to *Citrobacter freundii* and *E. coli* [34], potentiating synergism during polymicrobial infections. These examples suggest that the co-occurrence relationships we computationally detected may reflect genuine biological relationships. Further investigation of the interactions between different clusters of genera in the corrected sepsis network, together with expanding to future datasets, may yield valuable insights into the underlying biology of sepsis infections and ultimately inform treatment.

The presence of a densely connected cluster of oral colonisers may point to a potential reservoir of sepsis pathogens. This also suggests the possibility of opportunistic infections from the human microbiota and dysbioses that could affect disease severity. This hypothesis is in line with the reported changes in nasal microbiomes in septic individuals [35] and the associations of intestinal dysbiosis with increased susceptibility to sepsis [36]. If these hypotheses were true, microbiome profiles of patients might offer opportunities to assess a patient’s risk of developing sepsis prior to onset.

### The need to account for environmental contamination

Contamination from environmental sources poses one of the greatest challenges for metagenomic investigations of microbial communities, particularly in low biomass and clinical samples [20,37]. It is therefore crucial to discriminate between contaminants and biologically relevant taxa and to remove putative contaminants to protect against spurious signals.

The main premise behind SHAP Decontamination is that pathogens should occur at higher abundance in septic patients relative to healthy controls. This is because we expect most infections to be characterised by the proliferation of microorganisms [38,39] and, as such, true pathogenic genera should contribute to a higher predicted probability of sepsis at higher abundances. Consequently, the abundance of contaminant taxa would demonstrate a negative Spearman’s correlation with their corresponding SHAP values. This allows putative contaminant genera to be computationally detected and removed. Our results demonstrate the efficacy of our post-hoc contamination reduction technique called SHAP Decontamination in removing redundancy in the feature space while selectively retaining taxa involved in sepsis. It is likely that the taxa removed in this procedure would in principle include commensals and environmental contaminants introduced during sample collection or preparation. As such, application of this technique provides greater confidence that the polymicrobial signals we observed were not largely driven by contaminants.

We appreciate that a more rigorous evaluation of this technique, particularly with mock communities, will be required. As an alternative to our contamination reduction technique, statistical decontamination techniques identifying inverse relationships between the assigned abundance of taxa and sample DNA concentration [40,41] could be used. However, this method was not applicable for our study since the sample DNA concentrations in the datasets used were not reported.

### Potential for metagenomics-based diagnostics

Although we do not claim to have developed a model sufficiently robust for immediate diagnostic purposes, our results highlight the clear promise of metagenomics-informed diagnostic models, which have also been suggested by previous studies [22,42,43]. To put the high performance of our models in context, Mao et al. [9] reported that InSight, a model trained on vital signs of patients, had a diagnostic AUROC of 0.92 using Sepsis-2 as the ground truth. They also reported that the Modified Early Warning Score (MEWS), Sequential Organ Failure Assessment (SOFA) and SIRS had an AUROC of 0.76, 0.63 and 0.75 respectively. Additionally, a classifier trained on nasal metagenomes of septic and healthy samples had an AUROC of 0.89 with Sepsis-3 as the ground truth [35]. Notably, it is difficult to compare the performance of models trained with labels generated by different definitions of sepsis, which is also inherently a highly heterogeneous disease. Further, the discrepancies in model performance could be due to differences in the size of training and testing datasets. At the very least, our results suggest that the microbial component of sepsis alone contains sufficient information for the diagnosis of sepsis. A crucial next step will be to generate larger datasets, from more diverse sources, to allow the training of more robust and generalisable models for diagnostic or prognostic use.

### Limitations

We identified several limitations in our study. Firstly, metagenomic sequencing involves measurements of circulating free DNA and not of viable microorganisms in blood. As such, the detection of DNA from multiple taxa does not necessarily represent the true number or abundance of active taxa present. However, multiple studies have demonstrated high concordance of targeted [44] or shotgun metagenomic sequencing with culture [22,42,45]. This suggests some level of agreement between the presence of microbial cells and their DNA in blood. Additionally, given its higher sensitivity and throughput, metagenomic sequencing appears to be the best tool currently available for gaining insights into polymicrobial infections.

Though our results suggest the importance of multiple genera in delineating metagenomes of septic patients from that of healthy controls, the etiological contributions of these genera and their ecological relationships cannot be inferred. Such hypotheses must be confirmed experimentally. It is also important to keep in mind that the models presented in this study are not prognostic in nature, in that they were not trained to predict the onset or progression of sepsis. However, furthering our understanding of the microbial component of sepsis may prove useful in the development of better prognostic tools.

Some genera such as *Escherichia* and *Enterobacter* contain both biologically relevant genera and common sequencing contaminants. As such it is expected that a proportion of DNA molecules, and hence sequencing reads, may have come from contamination rather than microorganisms endogenous to blood. The abundance of these microorganisms, as detected by metagenomic approaches, may differ from the true abundance.

Additionally, *k*-mer based approaches may be less accurate for taxonomic classification compared to, for example, Bayesian sequence read-assignment methods [46]. As such, we used taxonomic assignments at the genus level which were shown to be, in general, more reliable than that at the species level [47]. We also appreciate that *k*-mer based classification approaches are significantly faster [48], which may provide clinically relevant turnaround times that are important in sepsis diagnostics.

Finally, we acknowledge the relatively small size of the datasets used in our analyses. As a result, the models presented in this study are not yet robust enough to be used in a clinical context. A larger and more diverse dataset is required to develop such models. This is to ensure that models can learn a more generalisable decision boundary for accurate sepsis diagnosis.

Irrespective of these limitations, our results nonetheless demonstrate the importance of considering the full polymicrobial component of sepsis and suggest that a metagenomics-based approach may provide biological and clinical insights supporting the future development of rapid diagnostic tools.

The advent of large-scale metagenomic sequencing of clinical samples offers new opportunities to better characterise the pathogens contributing to systemic infections, and unlike culture-based methods are not limited to organisms that are fast-growing or culturable. In this study, we demonstrate the promise of a metagenomics-based approach to sepsis. Our results provide evidence that septic infections should be considered as polymicrobial in nature, comprising multiple co-occurring pathogens indicative of disease. Our findings thus pave the way for more microbial-focused models of sepsis, with long run potential to inform early detection, clinical interventions and improve patient outcomes.

## Materials and Methods

### Datasets

Our primary analysis involved published shotgun metagenomic sequence data from the Karius study [22]. As detailed in this study, patients were diagnosed with sepsis if they presented with a temperature > 38°C or < 36°C, at least one other Systemic Inflammatory Response Syndrome (SIRS) criterion, and evidence of bacteraemia. Bacteraemia was confirmed via clinical microbiological testing performed within seven days after collection of the blood samples. The list of pathogens identified by such tests (which we refer to as ‘confirmed’ pathogens) can be found in Supplementary Table 5 of the Karius study, under the ‘definite’ adjudication. This included tissue, fluid and blood cultures, serology and nucleic acid testing. The clinical outcome of each patient was not reported in the original study. Seven of the 117 septic patients were found to have more than one ‘confirmed’ pathogen identified by microbiological testing (Supplementary Table 5; Karius study). According to the Karius study, healthy individuals were “screened for common health conditions including infectious diseases through a questionnaire and standard blood donor screening assays”. We believe this to be reasonable grounds for ruling out bloodstream infections in healthy patients (i.e. of non-septic origin).

### Data pre-processing

As described in the Karius study, input circulating free DNA was sequenced using NextSeq500 (75-cycle PCR, 1 x 75 nucleotides). Raw Illumina sequencing reads were demultiplexed by bcl2fastq (v2.17.1.14; default parameters) and quality trimmed using Trimmomatic (v0.32) [49] retaining reads with a quality (Q-score) above 20. Mapping and alignment were performed using Bowtie (v2.2.4) [50]. Human reads were identified by mapping to the human reference genome and removed prior to deposition in NCBI’s Sequence Read Archive (PRJNA507824).

For Grumaz-16 and Grumaz-19, *BBMap* (v38.79) [51] was used to trim adapter sequences, remove reads with a Q-score below 20 and remove reads mapping to the masked human hg19 reference (https://tinyurl.com/yya4xmrg). For the Gosiewski-17 dataset, we performed the same pre-processing steps as reported in the associated study [24]. Briefly, primers and adapters were removed using *Cutadapt* (v1.18) [52], paired reads merged using *ea-utils* (v1.1.2.537) [53], merged reads and forward unmerged *fastq* files concatenated, and reads with a Q-score below 20 removed using *BBMap*.

Taxonomic classification of all shotgun sequencing data was performed using *Kraken 2* (v2.0.9-beta; default parameters) [54] with the *maxikraken2_1903_140GB* database (https://tinyurl.com/y7zfg9kr). For the Gosiewski-17 dataset, *Kraken 2* with a *Kraken 2*-built *Silva* database was used instead of conventional 16S amplicon metagenomic classification methods [55]. Read assignments for all ‘confirmed’ bacterial pathogens using the *maxikraken2_1903_140GB* and *Kraken 2*-built *Silva* databases are shown in Fig. S2. While the relative number of reads assigned to each bacterial genus showed some inconsistencies, this hardly affected the classifier performance of septic and healthy patients (Fig. S3). This suggests that our model is fairly robust to heterogeneity which may be introduced by the classification step. For downstream analyses, we use the genera assignments based on the *Kraken 2*-built *Silva* database for the 16S Gosiewski-17 samples. Additionally, all unclassified reads were excluded from the analyses. The feature space obtained directly from *Kraken 2* taxonomic assignment is denoted by *Neat*.

Unexpectedly, for the Karius dataset, some reads were assigned to the genus *Homo* which was possibly due to misclassification. Mapping of all reads in the Karius sequencing data found just 873 bases with 96% identity to the masked human reference. Since human reads were already removed in the bioinformatic workflow of the Karius study, we did not perform an additional human read removal step to avoid introducing biases into the data.

### Model training, optimisation and evaluation

Classifiers were trained with a binary-logistic loss function and implemented using *XGBoost* API (v0.90) [56]. Model optimisation was performed using a randomised hyperparameter optimisation protocol [57] (1000 samples) implemented using *RandomizedSearchCV* in the *Scikit-learn* API (v0.23.1) [58]. The test error of each model was estimated using a nested, stratified, 10 x 10-fold cross-validation procedure. The best performing sets of hyperparameters that maximise the receiver operating characteristic curve (AUROC) of each model were determined and used for downstream analyses. The test error of each model was also estimated using a holdout test set after hyperparameter optimisation. For this procedure, precision, recall and AUPRC were used as performance metrics since they are more informative when used on imbalanced test sets [59].

### Model interpretation

To interpret models, each feature in a single sample was assigned a SHAP value, which corresponds to the change in a sample’s predicted probability score *(i.e.* probability of sepsis) when the feature is either present or absent. Using SHAP values therefore allows the decomposition of predicted probability scores for each sample into the sum of contributions from individual genera. The relative importance of each feature was inferred via its mean absolute SHAP value across all samples. A higher mean absolute SHAP value implies that the feature has a larger impact on the model predictions. SHAP values were computed using *TreeExplainer,* part of the *shap* library (v0.34.0) [26]. For every model, SHAP values were computed for the whole dataset by setting the *feature_pertubation* parameter to ‘interventional’.

### SHAP Decontamination

SHAP Decontamination was performed in two main steps. Firstly, genera that are not currently identified as known human pathogens were first removed. This selection was based on a study by Shaw *et al.* [60], who considered as a ‘human pathogen’ any microbial species for which there is evidence in the literature that it can cause infection in humans, sometimes in a single patient. Secondly, a classifier was optimised and trained on genera abundance *(Neat* feature spaces). SHAP values for model predictions on the dataset were then calculated. Genera with a negative Spearman’s correlation between their corresponding SHAP values and abundances were removed. Spearman’s correlations were calculated using *spearmanr* as part of the *SciPy* library (v1.4.1) [61]. A new classifier was then retrained using the previously optimised set of parameters but with this new reduced feature space. This process was repeated iteratively until the number of genera retained remained constant. The resultant feature space is denoted by *CR*.

To test the hypothesis that genera containing true pathogens are positively associated with sepsis, we inspected the SHAP values and read counts assigned to the genera corresponding to cases of each type of ‘confirmed’ infection *(e.g.* SHAP value/read count assigned to *Escherichia* for only *Escherichia-* positive samples) using the *Karius-Neat* feature space. The SHAP values were all at greater or equal to zero apart from a single sample which had a negative SHAP value for *Mycobacterium* (Fig. S4). The assigned read counts were non-zero except for one sample with a ‘confirmed’ fungal *Candida glabrata* infection reported (SRR8288759). These findings suggest that SHAP values can be used to identify experimentally identified pathogens.

### Simple Decontamination

We also employed a more direct, model-free contaminant removal technique (Simple Decontamination) that follows the same underlying premise of SHAP Decontamination. In this procedure, genera in the *Neat* feature space that were significantly *(p* < 0.05) more abundant in healthy controls than septic samples were considered contaminants and removed. The resultant feature space is denoted by *SD*.

### Microbial networks

Microbial co-occurrence networks were constructed using the *SparCC* algorithm [62], implemented in the *SpiecEasi* package (v1.1.0) [63] and visualised using *Igraph* (v1.2.5) [64]. *SparCC* was used to account for compositionality that could lead to spurious correlations. Separate networks were constructed for the genera assignments of septic and healthy metagenomes. To determine the microbial associations present exclusive to septic samples, a corrected sepsis network was produced. This network was constructed by subtracting all edges of the healthy network from the sepsis network. Only co-occurrence relationships where the *SparCC* correlations exceed 0.2 were retained. The *Karius-SD* feature space was used as input.

## Data Availability

All relevant source code and parsed datasets can be found on GitHub (https://github.com/cednotsed/Polymicrobial-Signature-of-Sepsis). The raw sequence data for each study can be found from NCBI SRA and the European Nucleotide Archive (ENA) repository with the accessions listed in Table 1.

