## Supplemental Materials for "Metagenomic evidence for a polymicrobial signature of sepsis" (from another version of this preprint: version 3, 2020-08-03)

### Supplementary Materials

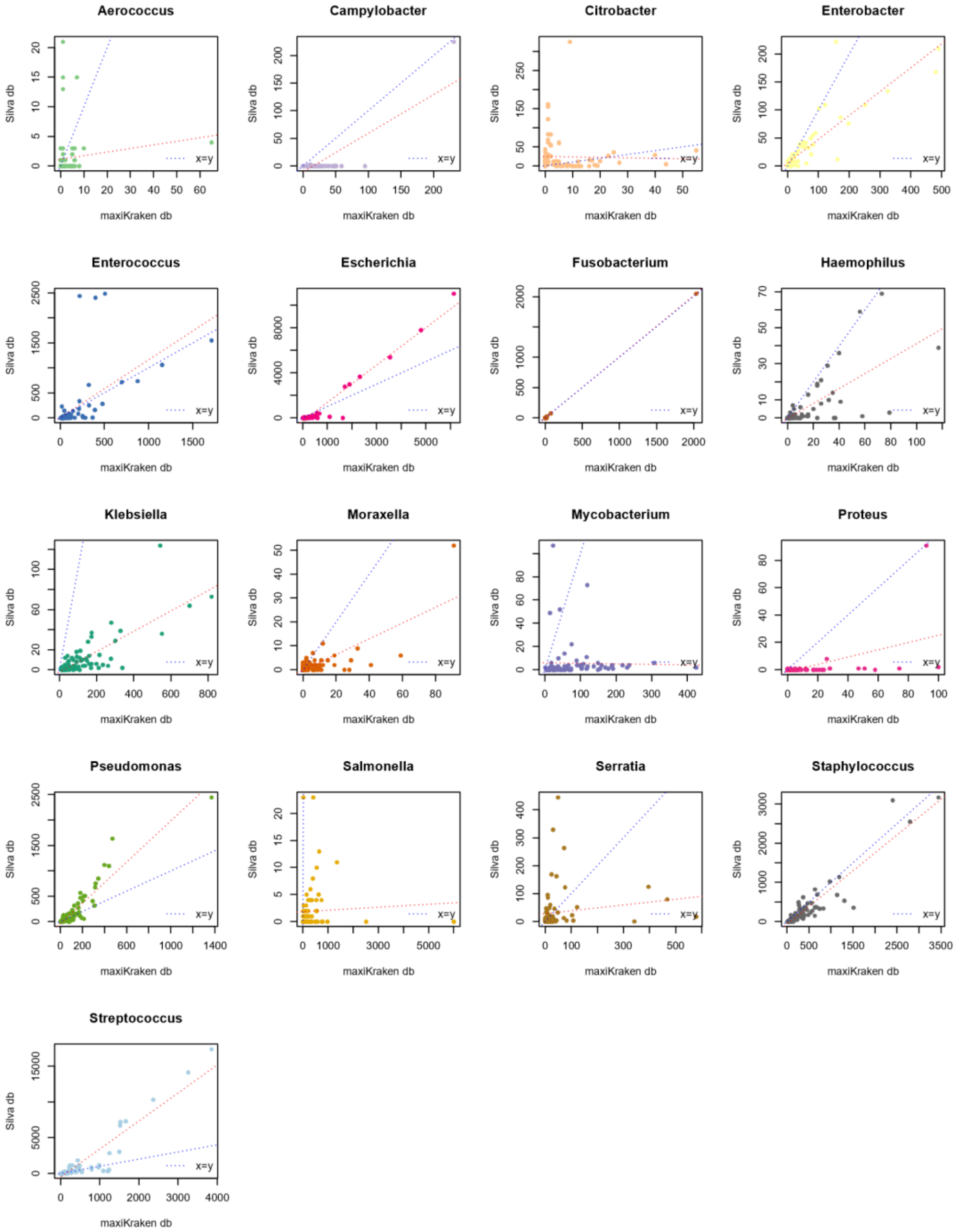


**Figure S1. Comparison of *Kraken 2* databases.** Genera abundance from taxonomic assignments using the *maxikraken2_1903_140GB* (denoted 'maxiKraken db') and the *Kraken 2*-built *Silva* database (denoted 'Silva db') for the Gosiewski-17 16S dataset. The 17 genera shown here correspond to those containing 'confirmed' pathogens in the Karius dataset excluding *Candida*, *Cryptococcus*, *Lymphocryptovirus*, *Simplexvirus* and *Cytomegalovirus*. The dashed red line provides the linear regression, with the blue dashed line providing the expectation under x=y.

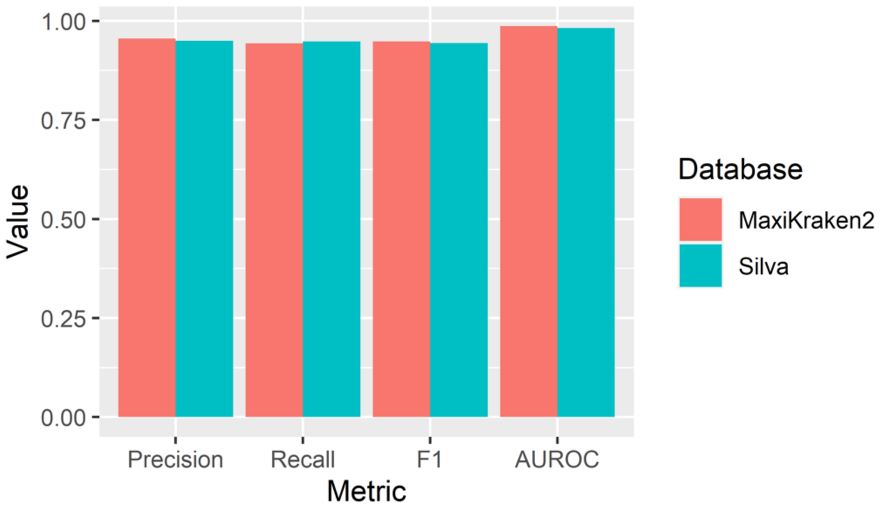


**Figure S2. Classification performance with different metagenomic reference databases.** Classifiers were trained on the *Pooled-Neat* feature spaces where the taxonomic assignment of the Gosiewski-17 16S dataset was either performed using the *maxikraken2_1903_140GB* (denoted 'MaxiKraken2') or the *Kraken 2*-built *Silva* database (denoted 'Silva'). Metrics were computed using the nested cross-validation protocol as described in the Methods section of the main text.

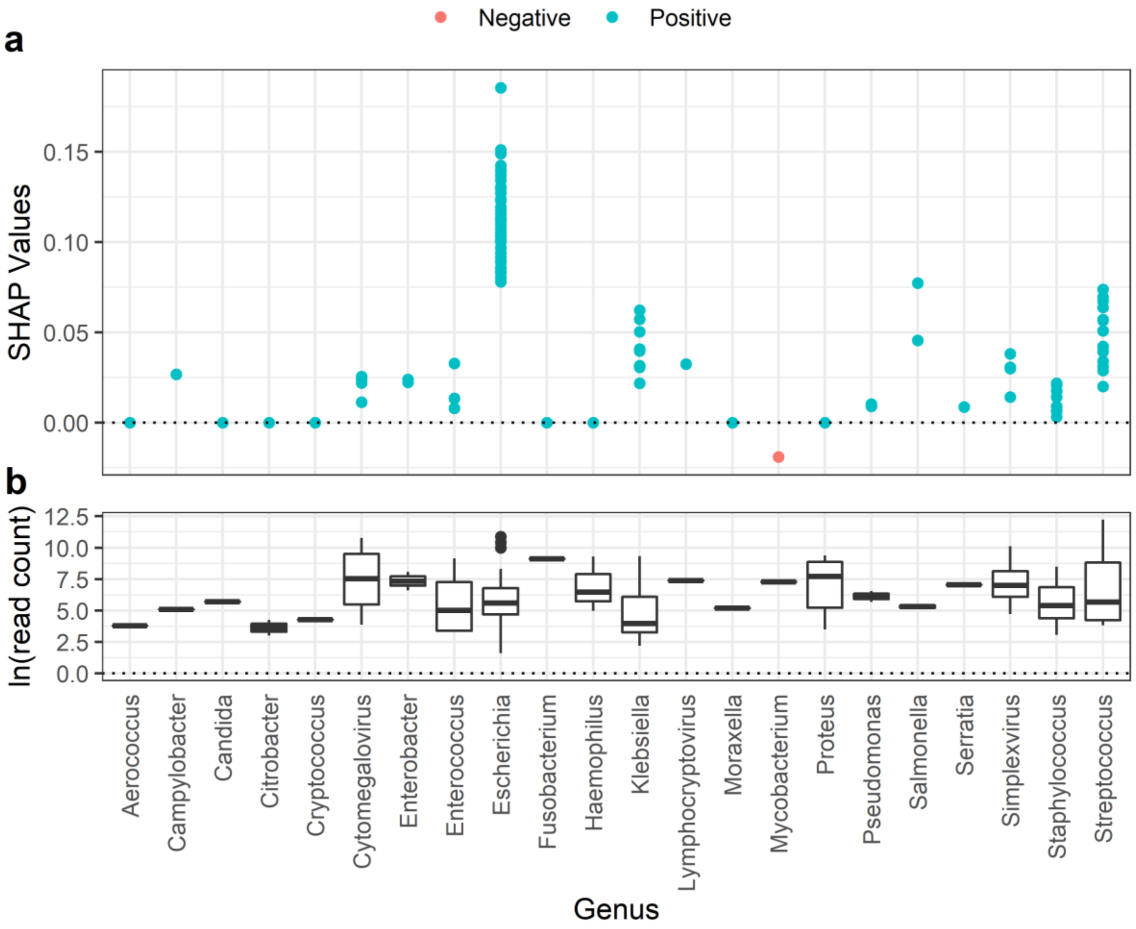


**Figure S3. SHAP values by infection type.** (a) Scatter plot of SHAP values (x-axis) of genera corresponding to sepsis samples for which a 'confirmed' infection was reported (y-axis). The SHAP values generated from the *Karius-Neat* model was used as input. Blue indicates values greater or equal to zero (denoted 'Positive') and red indicates values less than zero (denoted 'Negative'). (b) Abundance of genera for each type of 'confirmed infection' as measured by read counts on the natural logarithmic scale.

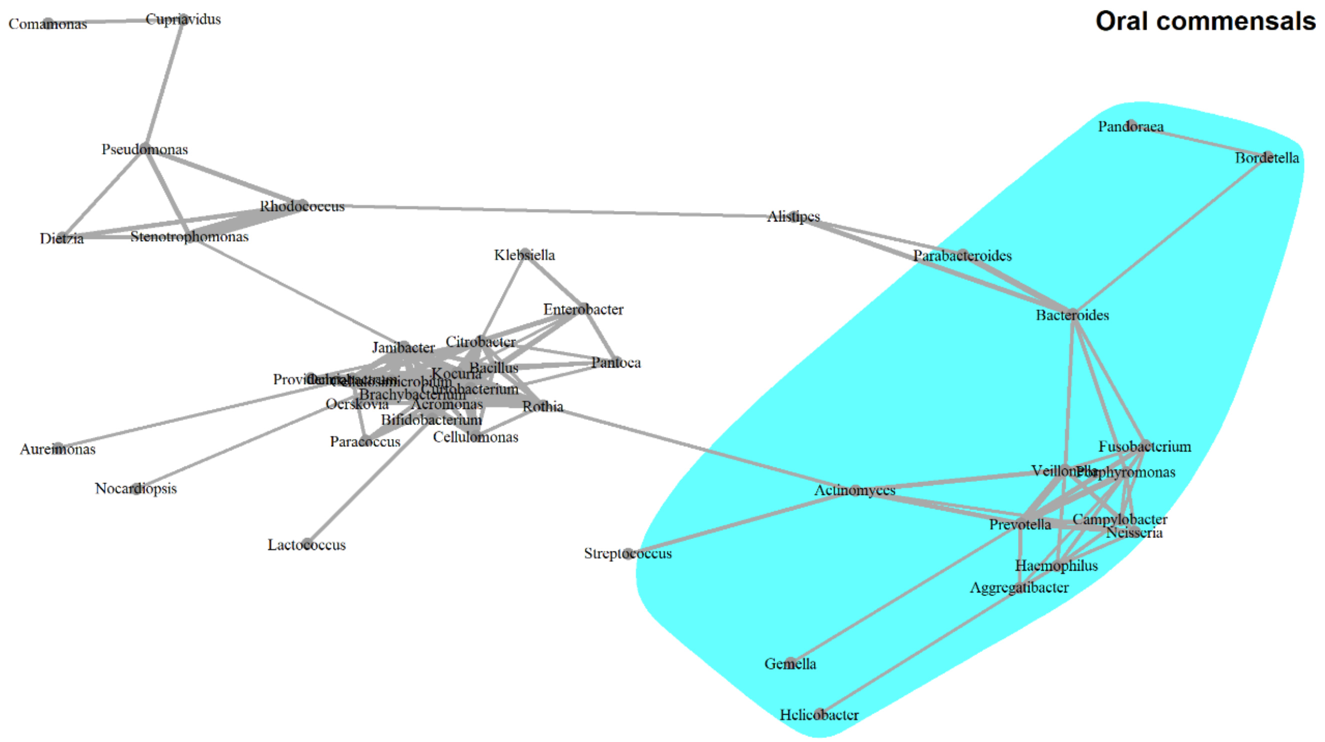


**Figure S4. Corrected microbial co-occurrence network for genera assigned in sepsis metagenomes.** Input data corresponds to the *Pooled-SD* feature space. The edges in this network represent those in the septic network that were not present in the healthy network. The widths of edges are weighted by the strength of the *SparCC* correlations. The layout of the graph was generated using the Fruchterman-Reingold algorithm.
